# Hydrogel-enhanced bioelectrochemical nitrate reduction for ammonium recovery from dilute nitrate via S*hewanella oneidensis* MR-1

**DOI:** 10.64898/2026.07.31.742018

**Authors:** Mamoru Oshiki, Yunjeong Choi, Ryuji Shinto, Satoshi Okabe

**Affiliations:** Division of Environmental Engineering, Faculty of Engineering, Hokkaido University, Japan

**Keywords:** bioelectrochemical nitrate reduction to ammonium, dilute nitrate, *Shewanella oneidensis* MR-1, hydrogel electrode

## Abstract

Bioelectrochemical reduction of dilute nitrate (NO₃⁻; sub-mM to low-mM range) to ammonium (NH₄⁺) offers a promising route toward circular nitrogen management from contaminated groundwater and environmental waters. However, on-site application of bioelectrochemical systems remains challenging due to low reduction rates and poor electron transfer efficiency of naturally formed biofilm electrodes. Here, we constructed a hydrogel biocathode by applying a carbon black/riboflavin/sodium alginate/cellulose hydrogel incorporating *Shewanella oneidensis* MR-1 cells to a graphite felt electrode via brush coating. The hydrogel electrode achieved NH₄⁺ production rates of 0.16–0.19 mol m⁻³ h⁻¹ without NO₂⁻ accumulation, and these rates were maintained without significant performance loss across three consecutive cycles with medium exchange over 1.5 days of total operation. The hydrogel electrode increased the current density by more than 5-fold compared with a conventional *S. oneidensis* biofilm electrode, indicating enhanced electron transfer efficiency per unit biomass, which directly contributed to the high NH₄⁺ production rates. The electricity consumption for NH₄⁺ production of 1.68–2.39 × 10² kJ g-N⁻¹ was substantially lower than that of metal catalyst systems at comparable NO₃⁻ concentrations (typically, >10^4^ kJ g-N⁻¹). These findings demonstrate that the hydrogel electrode design represents an energy-efficient, and readily fabricated platform for bioelectrochemical NH₄⁺ production from dilute NO₃⁻.

## 1. Introduction

The Haber–Bosch process for industrial nitrogen fixation is a cornerstone technology underpinning modern agriculture, consuming approximately 1–2% of global primary energy [1] while producing roughly 160 Tg-N of NH₄⁺ annually [2,3] However, more than half of the fixed NH₄⁺ is not taken up by crops and instead leaches into the environment as NO₃⁻ [2,4], contaminating groundwater and surface waters. To mitigate NO₃⁻ pollution, biological nitrogen removal (denitrification and anaerobic ammonium oxidation) technologies have been developed and widely deployed. However, these processes are energy-intensive; for example, wastewater treatment accounts for approximately 1% of national electricity consumption [5,6]. This represents a fundamentally unidirectional system in which enormous amounts of energy are first expended to fix nitrogen (*i.e*., the Haber–Bosch process), only for comparably large amounts of energy to then be consumed to return it to the atmosphere as N_2_ gas. The current nitrogen cycle thus depends on an unsustainable “fix → dissipate → remove” structure, and there is a growing need to transition toward “circular nitrogen management” — a paradigm in which dissipated NO₃⁻ is not merely removed but recovered and recycled as a nitrogen resource (NH₄⁺) [7]. Of particular concern is the widespread occurrence of relatively low-concentration NO₃⁻ (dilute NO₃⁻) in the range of hundreds of µM to several mM in environmental waters [8–10], underscoring the urgent need to develop NO₃⁻ reduction and NH₄⁺ recovery technologies that can operate efficiently even under such dilute conditions.

As an emerging approach for NH₄⁺ recovery from NO₃⁻, electrochemical reduction using metal catalysts has seen particularly rapid development in recent years [9,11]. However, metal catalyst systems generally suffer from a pronounced decrease in reaction efficiency as substrate concentration decreases [10,12]. As noted above, NO₃⁻ concentrations in environmental waters are typically on the order of ∼1 mM, conditions that are inherently unfavorable for metal catalysts. On the other hand, enzymes have evolved highly specific substrate-recognition mechanisms. NO₃⁻ is sequentially reduced to NH₄⁺ via nitrate reductases (Nap or Nar) and NH_4_^+^–forming nitrite reductase (Nrf) [13], and the apparent affinity constants of bacterial Nap for NO_3_^−^ and Nrf for NO_2_^−^ are typically within the submillimolar range [13,14]. These high apparent substrate affinities enable biocatalysts to efficiently reduce dilute NO_3_^−^ in environmental waters. A few microbial electrosynthesis (MES) systems for NO₃⁻ reduction and NH₄⁺ recovery using biocatalysts have been reported to date [15–18]. However, most of these studies rely on biofilms formed by the natural attachment and proliferation of microorganisms on the cathode surface, and such naturally formed biofilms are physically fragile. In the field of bioelectrochemical systems, various approaches to modifying cathode materials to improve electrode performance have been reported. In particular, Yang et al. [19] recently demonstrated in a microbial fuel cell (MFC) anode system that incorporating a hydrogel matrix for cell immobilization as the electrode material substantially improved current density and reduced charge transfer resistance. However, no study has yet applied such hydrogel electrodes to the cathode of an MES system for NO₃⁻ reduction and NH₄⁺ recovery.

In this study, *Shewanella oneidensis* MR-1 — which possesses both an extracellular electron transfer (EET) system capable of direct electron exchange with electrodes [20] and the metabolic potential for dissimilatory nitrate reduction to ammonium (DNRA) [21] — was employed as the biocatalyst, and an MES was constructed by applying a hydrogel electrode composed of carbon black (CB)/riboflavin (RF)/sodium alginate (SA)/cellurose (Cel) as the cathode [19]. Using this MES, the NO_3_^−^ reduction and NH₄⁺ production performance under dilute NO₃⁻ conditions was evaluated. The electrochemical properties of the hydrogel electrode were further characterized by electrochemical analyses. This study demonstrates that the use of a hydrogel electrode enables high and stable bioelectrochemical NO₃⁻ reduction and NH₄⁺ recovery performance, thereby expanding the application of MES technology.

## 2. Material and methods

### 2.1 Cultivation of *S. oneidensis* MR-1

*S. oneidensis* MR-1 (JCM31522) was inoculated into 25 mL of 1 × Luria-Bertani medium (Nacalai Tesque, Kyoto, Japan) and cultivated aerobically at 30°C and 90 rpm for 2 days. After cultivation, the culture was transferred to 50 mL centrifuge tubes and centrifuged at 8,000 × g for 10 min at 4°C to pellet the cells. The supernatant was removed by pipetting, and the cells were washed three times with 1 × phosphate-buffered saline (PBS) to thoroughly remove residual medium components. The resulting *S. oneidensis* cell suspension was used for cathode fabrication as described below.

### 2.2 Fabrication of biocathode

In this study, two types of biocathode were prepared: (1) a cathode in which *S. oneidensis* cells were encapsulated in a CB/RF/SA/Cel hydrogel, and (2) a biofilm cathode on which *S. oneidensi*s was allowed to form a biofilm autonomously.

#### 2.2.1 Fabrication of hydrogel electrode

The hydrogel bioink was prepared following the procedure reported by Yang et al. [19]. Briefly, 0.4 g of carbon black (CB) Vulcan XC72R (Cabot Corporation; pore size 20–50 nm) was added to 20 mL of deionized water in a 50 mL centrifuge tube and dispersed by ultrasonication for 10–20 min using an ultrasonic bath (US-107, SND Co., Ltd., Nagano, Japan). Riboflavin (RF; Fujifilm Wako Pure Chemical Corporation, Osaka, Japan) was then added to a final concentration of 10 mM, and the mixture was rotated in the dark at room temperature (<5 rpm) for 24 h. The tube was centrifuged (8,000 × *g*, 10 min), and the pellet was collected as the CB/RF composite. To the CB/RF composite, 0.04 g of sodium alginate (SA; viscosity 100–150 cP; Fujifilm Wako Pure Chemical Corporation) and 0.1 g of α-cellulose (Cel; Sigma Life Science, St. Louis, MO, USA) were added and suspended in 2.5 mL of deionized water to prepare a CB/RF/SA/Cel mixture. The bioink was prepared by mixing 2.5 mL of the CB/RF/SA/Cel mixture with 2.5 mL of *S. oneidensis* cell suspension in PBS.

Prior to bioink application, graphite felt electrodes (2 × 7.5 × 1 cm; AS ONE Co., Japan) were pre-treated by immersion in 1 M HNO₃ followed by ultrasonication for 5–10 min (US-107, SND Co., Ltd.). The electrodes were then rinsed repeatedly with deionized water until the rinse solution reached neutral pH. The bioink was applied to the pre-treated graphite felt by brush coating at a loading of 167 µg cm⁻³ (mass of bioink per unit electrode volume): 1.5 mL was applied to one face of the 2 × 7.5 cm surface, and 0.5 mL was applied to each face of the 7.5 × 1 cm surfaces. After application, the graphite felt was immersed in 0.5 M CaCl₂ solution at room temperature for 2 min to crosslink the sodium alginate and form the hydrogel.

#### 2.2.2 Fabrication of biofilm electrode

A *S. oneidensis* biofilm was formed on the electrode surface by electrochemical cultivation in the anode chamber of an H-type glass reactor, using lactate as the electron donor and the electrode as the electron acceptor [22]. A glass H-type reactor (anode and cathode chamber volume: 300 mL each) equipped with a Nafion 117 proton exchange membrane (PEM) was used [23]. The anode solution (composition: Table S1) and cathode solution (Table S4) were dispensed at 300 mL each. After dispensing, the anode solution was sparged aseptically with N₂ gas for at least 15 min. Graphite felt electrodes connected with titanium wire were used as counter electrodes, and an Ag/AgCl electrode (RE-1A, EC Frontier Co., Ltd., Kyoto, Japan) was used as the reference electrode. After inoculation of *S. oneidensis* into the anode chamber, the anode potential was poised at +0.2 V vs. Ag/AgCl using a potentiostat (HA-151B, Hokuto Denko Corporation, Tokyo, Japan) and the reactor was operated in MFC mode under anaerobic conditions. The time course of current during biofilm formation is shown in Fig. S1. After 72 h, the graphite felt electrode was removed from the anode chamber and gently rinsed with 1× PBS to remove loosely attached components. The resulting biofilm electrode was used in subsequent experiments.

### 2.3 Bioelectrochemical NO_3_^−^ reduction to NH_4_^+^

The same reactor configuration and operational conditions were applied for both the hydrogel and the biofilm electrodes. A glass H-type reactor (anode and cathode chamber volume: 300 mL each) equipped with a Nafion 117 PEM was used. The compositions of the cathode and anode solutions are provided in Tables S5 and S7, respectively. Prior to each experiment, the cathode chamber was sparged with Ar gas for 10 min, and anoxic conditions were subsequently maintained by sealing the reactor. The hydrogel or biofilm electrodes (2 sheets) were placed in the cathode chamber, and a graphite felt connected with titanium wire was inserted into the anode chamber. An Ag/AgCl electrode was used as the reference electrode. The cathode potential was controlled at −0.7 V vs. Ag/AgCl using a potentiostat (HA-151B, Hokuto Denko Corporation). All experiments were conducted at a constant temperature of 25°C.

Experiments using the hydrogel electrode were performed in triplicate using three independently fabricated reactors (n = 3). When NO₃⁻ in the cathode solution was depleted, the solution was replaced with fresh cathode solution containing NO₃⁻, and the operation was repeated for a total of three cycles (Cycles 1–3). Experiments using the biofilm electrode were conducted following the same procedure (Cycles 1–2). Control experiments were performed under (i) open-circuit conditions without applied potential and (ii) conditions using a hydrogel electrode without *S. oneidensis* cells. Liquid samples were withdrawn from the cathode chamber during operation, filtered through a 0.45 µm membrane filter, and subjected to chemical analysis as described below.

### 2.4 Chemical analysis

The concentrations of NO₃⁻, NO₂⁻, and NH₄⁺ were determined colorimetrically or fluorometrically using a multimode microplate reader (Nivo™, PerkinElmer, Waltham, MA, USA). The NO₃⁻ concentration was determined by either the brucine sulfuric acid method or the hydrazine-copper reduction method. For the brucine sulfuric acid method, 16 µL of NaCl solution, 80 µL of H₂SO₄ solution, and 4 µL of brucine solution were added to 80 µL of liquid sample. After heating at 95°C for 20 min, 20 µL of distilled water was added and the mixture was cooled to room temperature. Absorbance was measured at 405 nm [24]. For the hydrazine-copper reduction method, 105 µL of distilled water, 30 µL of catalyst solution (CuSO₄·5H₂O 35.4 mg/L, ZnSO₄·7H₂O 1.44 g/L), 30 µL of 1 M NaOH solution, and 30 µL of hydrazine solution were added sequentially to 5 µL of liquid sample in a 96-well plate and mixed. After standing at room temperature for 15 min, 32 µL of naphthylethylenediamine solution was added. Absorbance was measured at 540 nm [25]. The NO₂⁻ concentration was determined by the naphthylethylenediamine colorimetric method. Briefly, 195 µL of distilled water was added to 5 µL of liquid sample in a 96-well plate to give a total volume of 200 µL, followed by addition of 8 µL of naphthylethylenediamine solution. After standing at room temperature for 5 min, absorbance was measured at 540 nm [26]. The NH₄⁺ concentration was determined by the OPA (ortho-phthaldialdehyde) fluorometric method [27]. Briefly, 200 µL of OPA reagent (1×) was added to 10 µL of liquid sample in a 96-well plate. After incubation at room temperature for 20 min in the dark, fluorescence was measured at an excitation wavelength of 355 nm and an emission wavelength of 460 nm.

The ¹⁵N/¹⁴N ratio of NH₄⁺ was determined by ¹H-NMR spectroscopy [28]. Liquid samples (0.55 mL) were transferred into 5 mm NMR tubes, mixed with DMSO-*d*₆ as a deuterated solvent at a ratio of 9:1 (v/v), and supplemented with maleic acid as an internal standard at a final concentration of 1.3 mM. The pH of each sample was adjusted to approximately 3–4 by addition of sulfuric acid solution. ¹H-NMR spectra were recorded on a JEOL ECA600 spectrometer (JEOL Ltd., Tokyo, Japan) operating at 600 MHz. The ¹⁴NH₄⁺ and ¹⁵NH₄⁺ signals were distinguished based on their characteristic splitting patterns: ¹⁴NH₄⁺ (spin quantum number *I* = 1) appears as a 1:1:1 triplet, whereas ¹⁵NH₄⁺ (spin quantum number *I* = 1/2) appears as a 1:1 doublet. The ¹⁵N fraction of NH₄⁺ was calculated from the integrated peak areas of the respective signals according to the following equation:

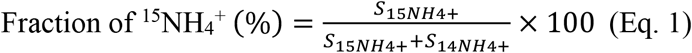

where *S_14NH4+_* and *S_15NH4+_* denote the integrated areas of the triplet peak derived from ¹⁴NH₄⁺ and the doublet peak derived from ¹⁵NH₄⁺, respectively.

### 2.5 Electrochemical analysis

Electrochemical measurements were performed using an electrochemical analyzer (ALS 700E, BAS Inc., Tokyo, Japan) in a three-electrode configuration, in which the cathode served as the working electrode, the anode as the counter electrode, and an Ag/AgCl electrode as the reference electrode [22,23]. Linear sweep voltammetry (LSV) was conducted over a potential range of −0.8 to −0.15 V vs. Ag/AgCl at a scan rate of 1 mV/s. Cyclic voltammetry (CV) was performed over a potential range of −0.8 to +0.2 V vs. Ag/AgCl at a scan rate of 50 mV/s. Electrochemical impedance spectroscopy (EIS) was conducted at a cathode potential of −0.7 V vs. Ag/AgCl over a frequency range of 10⁵ to 0.01 Hz with a sinusoidal perturbation amplitude of 10 mV.

### 2.6 Quantification of *S. oneidensis* cell-associated biomass on electrodes

The hydrogel and biofilm electrodes were immersed in 0.2 N NaOH solution and heated at 96°C for 20 min to lyse the *S. oneidensis* cells in the hydrogel electrode and the cells attached to the biofilm electrode. The protein concentration in the lysate was then quantified using the NanoOrange Protein Quantitation Kit (Thermo Fisher Scientific, Waltham, MA, USA) according to the manufacturer’s protocol. A standard curve was constructed using bovine serum albumin (BSA).

### 2.7 Calculation of performance indices

Faradaic efficiency (FE) was calculated according to the following equation:

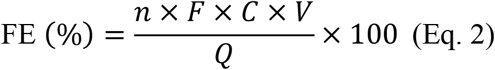

where *n* is the number of electrons required per mole of NO₃⁻ reduced to NH₄⁺ (8 *e⁻* mol⁻¹), *F* is the Faradaic constant (96,485 C mol⁻¹), *C* is the NH₄⁺ concentration (mol L⁻¹), *V* is the volume of the cathode solution (L), and *Q* is the total charge transferred (C).

The NH₄⁺ production rate was calculated according to the following equation:

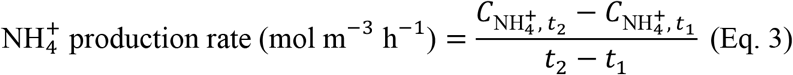

where 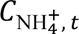 is the NH₄⁺ concentration at time *t* (mol m⁻³), *t*₁ = 3 h, and *t*₂ = 12 h. The nitrogen mass balance was calculated according to the following equation:

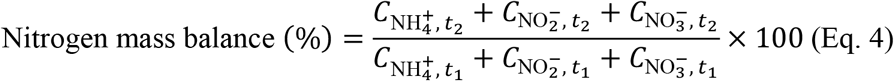

where 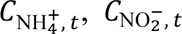 and 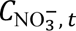 are the concentrations of NH₄⁺, NO₂⁻, and NO₃⁻ at time *t* (mM), respectively; *t*₁ = 0 h and *t*₂ = 12 h.

The selectivity of NO₃⁻ reduction to NH₄⁺ was calculated according to the following equation:

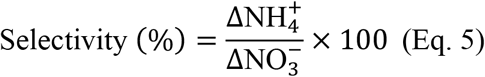

where 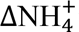 is the increase in NH₄⁺ concentration from 0 to 12 h (mM) and 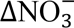 is the decrease in NO₃⁻ concentration from 0 to 12 h (mM).

The electricity consumption (EC; kJ g-N⁻¹) was calculated by dividing the total electrical energy input by the mass of NO₃⁻-N removed (Eq. 3) [29]:

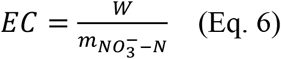

where *EC* is the electricity consumption, *w*is the electrical energy input, and 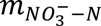 is the mass of NO₃⁻-N removed during the reaction.

The electrical energy input was calculated from the cell voltage and current (Eq. 4):

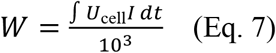

where *U*_cell_ is the potential difference between the anode and cathode chambers, *I* is the current, and *t* is the operation time. The factor 10^3^ was used to convert joules to kilojoules. The final EC unit was expressed as kJ g-NO₃⁻-N⁻¹.

### 2.8 Statistical analysis

Differences in NH₄⁺ production rate, current, FE, and EC among the three operational cycles were evaluated using the Friedman test as a non-parametric test for related samples, followed by pairwise post-hoc comparisons using paired *t*-tests. The resulting *p*-values were corrected for multiple comparisons using the Holm method. All statistical analyses were performed in Python using the *scipy.stats* and *statsmodels* packages. A significance level of *α* = 0.05 was applied.

## 3. Results

### 3.1 Bioelectrochemical NO_3_^−^ reduction to NH_4_^+^ with *S. oneidensis* hydrogel

A hydrogel electrode containing *S. oneidensis* was formed on a graphite felt electrode and operated as a biocathode at −0.7 V vs. Ag/AgCl for bioelectrochemical NO₃⁻ reduction (Fig. 1a). In Cycle 1, NO₃⁻ concentration decreased over 12 h, with concomitant accumulation of NH₄⁺ (Fig. 1b). No significant accumulation of NO₂⁻ was observed. In Cycle 1, ¹⁵NO₃⁻ (^15^NaNO_3_, >98% ^15^N/^14^N) (Shoko Science, Japan) was supplied as a tracer, and the ¹⁵N/¹⁴N ratio of NH₄⁺ was measured at 12 h, yielding a value of 81 ± 1.9% (mean ± SD, *n* = 3). After 12 h of operation, the medium was replaced with fresh medium (containing ¹⁴NO₃⁻ instead of ¹⁵NO₃⁻), and the operation was repeated (Cycles 2 and 3). In all cycles, the same trend of NO₃⁻ decrease and NH₄⁺ accumulation was reproducibly observed. Regarding the nitrogen mass balance, values of 118 ± 3.1%, 160 ± 3.6%, and 128 ± 25% were obtained for Cycles 1, 2, and 3, respectively, and the selectivity was 100 ± 3.4%, 127 ± 18%, and 99 ± 30% for Cycles 1, 2, and 3, respectively. As negative controls, the experiments were repeated under (i) open-circuit conditions without applied voltage and (ii) conditions in which a hydrogel electrode without *S. oneidensis* inoculation was subjected to the same applied potential; NO₃⁻ consumption was not observed under either condition.

**Fig. 1.**
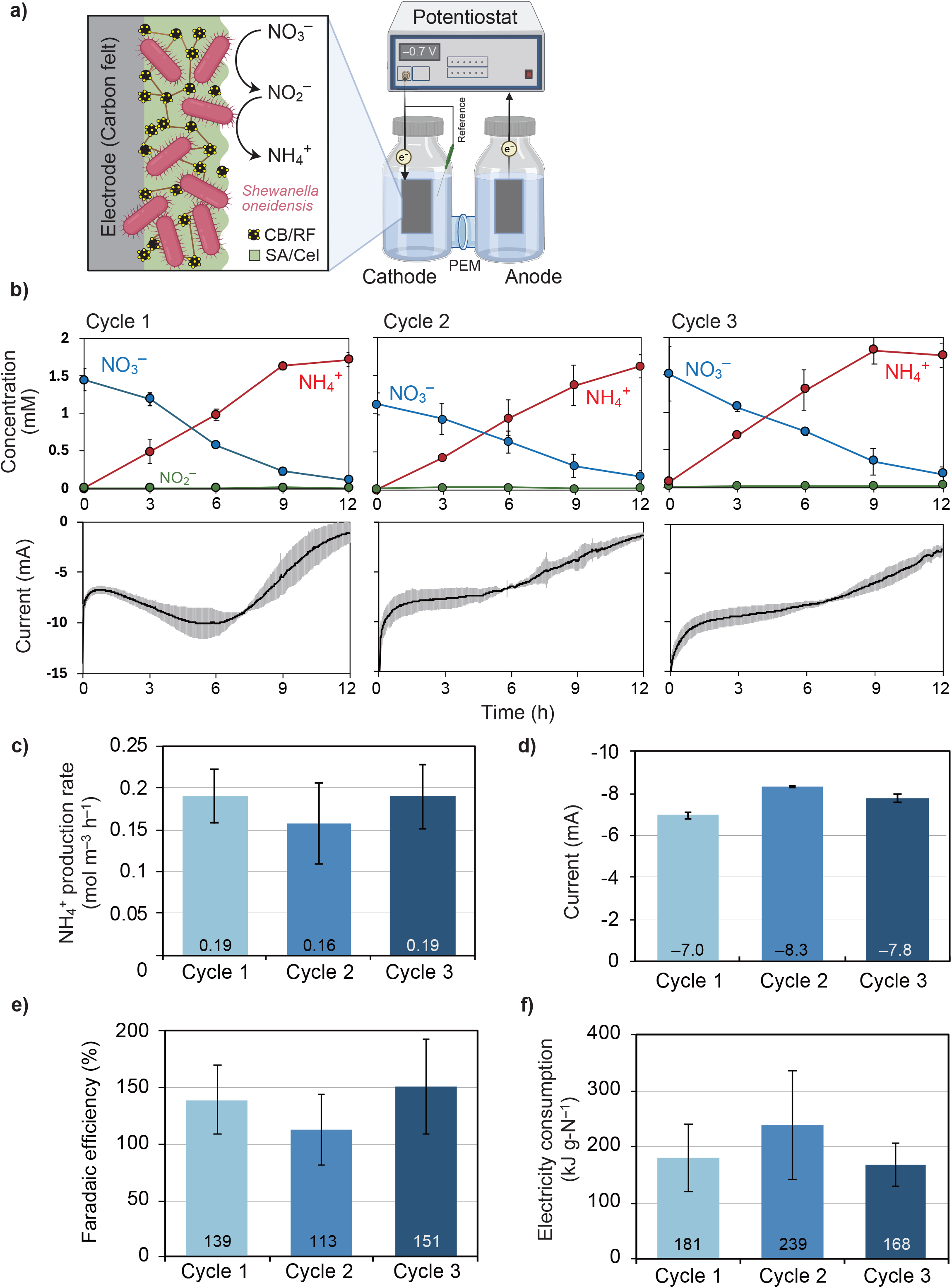
Bioelectrochemical NO_3_⁻ reduction to NH_4_⁺ using a *S. oneidensis* hydrogel electrode. (a) Schematic illustration of the experimental setup. *S. oneidensis* MR-1 immobilized in a CB/RF/SA/Cel hydrogel was applied to a carbon felt electrode as a biocathode and operated at a constant potential of −0.7 V vs. Ag/AgCl. **(b)** Time courses of NO_3_⁻, NO_2_⁻, and NH_4_⁺ concentrations (upper panels) and current (lower panels) during each cycle. Error bars indicate standard deviations of three independent replicates (*n* = 3). **(c)** NH_4_⁺ production rate, **(d)** current (average over 3–12 h), **(e)** Faradaic efficiency (FE), and **(f)** electricity consumption (EC) for each cycle. Bars represent means of *n* = 3; error bars indicate standard deviations.

The NH₄⁺ production rate, current, FE, and EC were calculated for Cycles 1 through 3 (Fig. 1c–f). The NH₄⁺ production rate was 0.19 ± 0.03, 0.16 ± 0.05, and 0.19 ± 0.04 mol m⁻³ h⁻¹ for Cycles 1, 2, and 3, respectively; the current was −7.0 ± 0.18, −8.3 ± 0.07, and −7.8 ± 0.21 mA; the FE was 139 ± 31%, 113 ± 32%, and 151 ± 43%; and the EC was 181 ± 61, 239 ± 96, and 168 ± 38 kJ g-N⁻¹. No statistically significant differences were observed among cycles for any of these parameters, demonstrating stable NO₃⁻ reduction and NH₄⁺ production performance over a total of three cycles.

For comparison with the hydrogel electrode, an experiment was conducted in which a *S. oneidensis* biofilm was directly formed on the electrode surface without hydrogel (Fig. S2). Although NO₃⁻ reduction and NH₄⁺ production occurred, their rates were lower than those of the hydrogel system; *i.e.,* NH₄⁺ production rate: 0.014 mol m⁻³ h⁻¹. Also, the current (−1.28 mA) was >5 folds lower than those found on the hydrogel system. When the medium was replaced after 96 h and operation was resumed in Cycle 2, both the NH₄⁺ production rate and current declined further; *i.e*., 0.0023 mol m⁻³ h⁻¹ and −0.47 mA, respectively.

### 3.2 Electrochemical characterization of the cathode electrode

The electrochemical properties of the hydrogel electrode (0 h of incubation) were evaluated by LSV (Fig. 2a). The hydrogel electrode containing *S. oneidensis* exhibited a distinct reduction peak at −0.648 V. In contrast, the hydrogel without *Shewanella* cells showed a peak at −0.529 V, and no distinct peak was observed for the *Shewanella* biofilm electrode. The hydrogel electrode containing *S. oneidensis* showed a 6.1-fold increase in current density compared to the biofilm electrode: −0.888 and −0.145 mA cm⁻³ at −0.648 V and −0.8 V, respectively. In CV measurements, the *S. oneidensis* hydrogel electrode exhibited a markedly larger CV area compared to the *S. oneidensis* biofilm electrode (Fig. 2b). However, no distinct redox peaks were observed for either electrode in the CV curves.

**Fig. 2.**
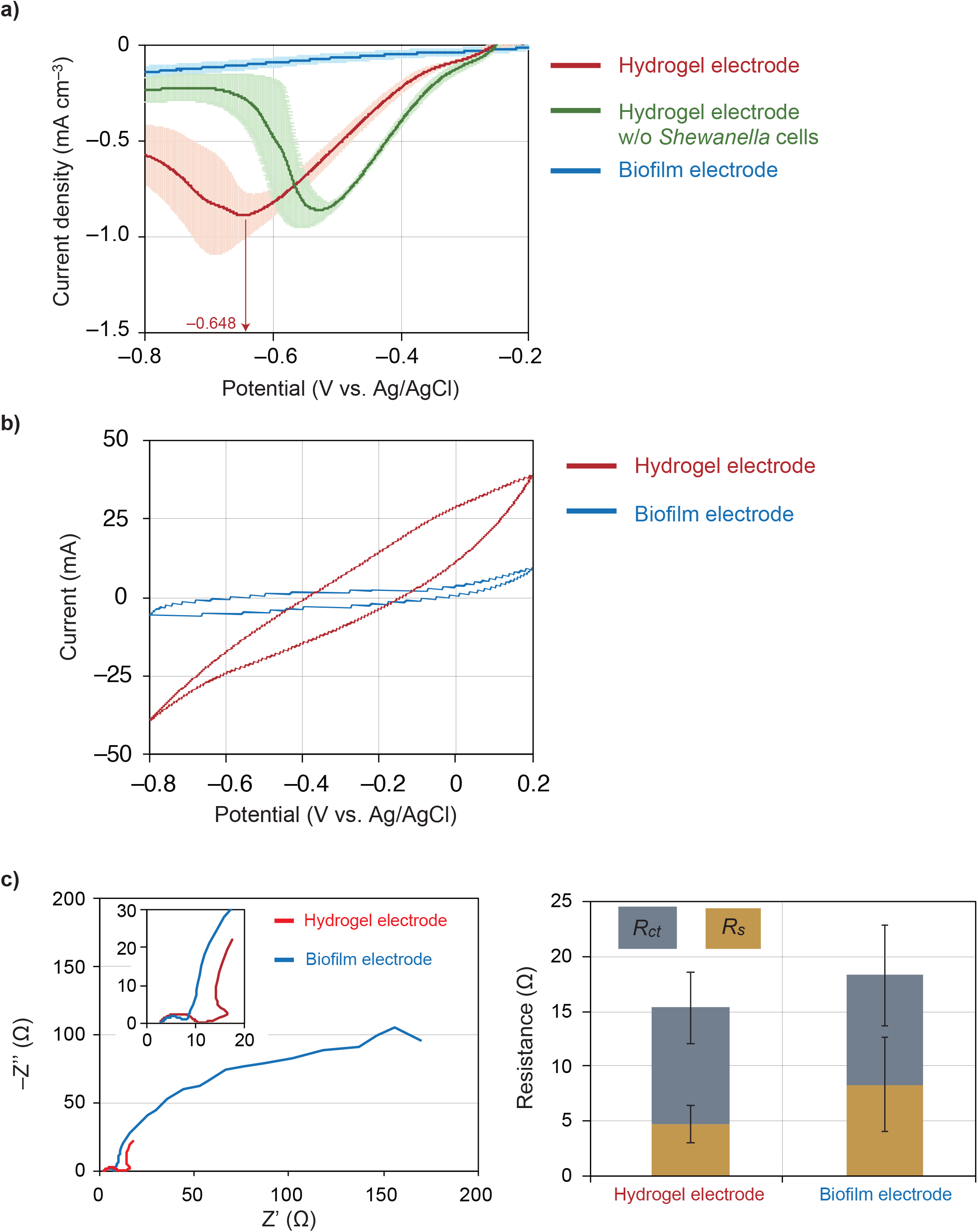
Electrochemical characterization of the hydrogel electrode. **(a)** Linear sweep voltammetry (LSV) curves of the CB/RF/SA/Cel hydrogel electrode (red), the hydrogel electrode without *Shewanella* cells (green), and the *Shewanella* biofilm electrode (blue). Scan rate: 1 mV/s. **(b)** Cyclic voltammetry (CV) curves of the hydrogel electrode (red) and the *Shewanella* biofilm electrode (blue). Scan rate: 50 mV/s. **(c)** Nyquist plots from electrochemical impedance spectroscopy (EIS) (left) and charge transfer resistance (*R_ct_*) and solution resistance (*R_s_*) derived from equivalent circuit fitting (right) for the hydrogel electrode (red) and the biofilm electrode (blue).

Electrode resistance was evaluated by EIS (Fig. 2c). Nyquist plot analysis and equivalent circuit fitting revealed no significant difference in charge transfer resistance (*R_ct_*) or solution resistance (*R_s_*) between the hydrogel and biofilm electrode: *R_ct_*: 10.6 ± 3.23 and 9.95 ± 4.56 Ω; *R_s_*: 8.43 ± 4.26 and 4.83 ± 1.71 Ω, respectively.

To evaluate the biomass on the electrodes, bacterial proteins were extracted by NaOH treatment from the hydrogel electrode (Fig. 1) and the biofilm electrode (Fig. S2) at the start of Cycle 1. No significant difference in protein amounts was observed between those two electrodes: 2.3 ± 0.6 and 2.7 ± 0.2 µg-protein cm⁻³ electrode, respectively.

### 3.3 Performance comparison with previously reported NO_3_^−^ reduction system

The volumetric NH₄⁺ production rate and FE achieved in the present study were plotted with the values reported in the literature for bioelectrochemical NO₃⁻ reduction systems (Fig. 3a). Both the NH₄⁺ production rate (0.16–0.19 mol m⁻³ h⁻¹) and FE (113–151%) achieved in this study (*S. oneidensis* hydrogel) exceeded those reported for *S. oneidensis* biofilm [15], *S. oneidensis* biofilm & FeS [16], and *Geobacter* sp. biofilm [17]. Furthermore, a comparison with metal catalyst-based electrochemical NO₃⁻ reduction systems [10] was conducted (Fig. 3b). While metal catalysts have been reported to require EC values of nearly 10^3^ kJ g–N^−1^ or higher at mM-level NO₃⁻ concentrations, the *S. oneidensis* hydrogel system in the present study achieved an EC of 1.68–2.39 × 10² kJ g-N⁻¹ under comparable substrate concentration conditions.

**Fig. 3.**
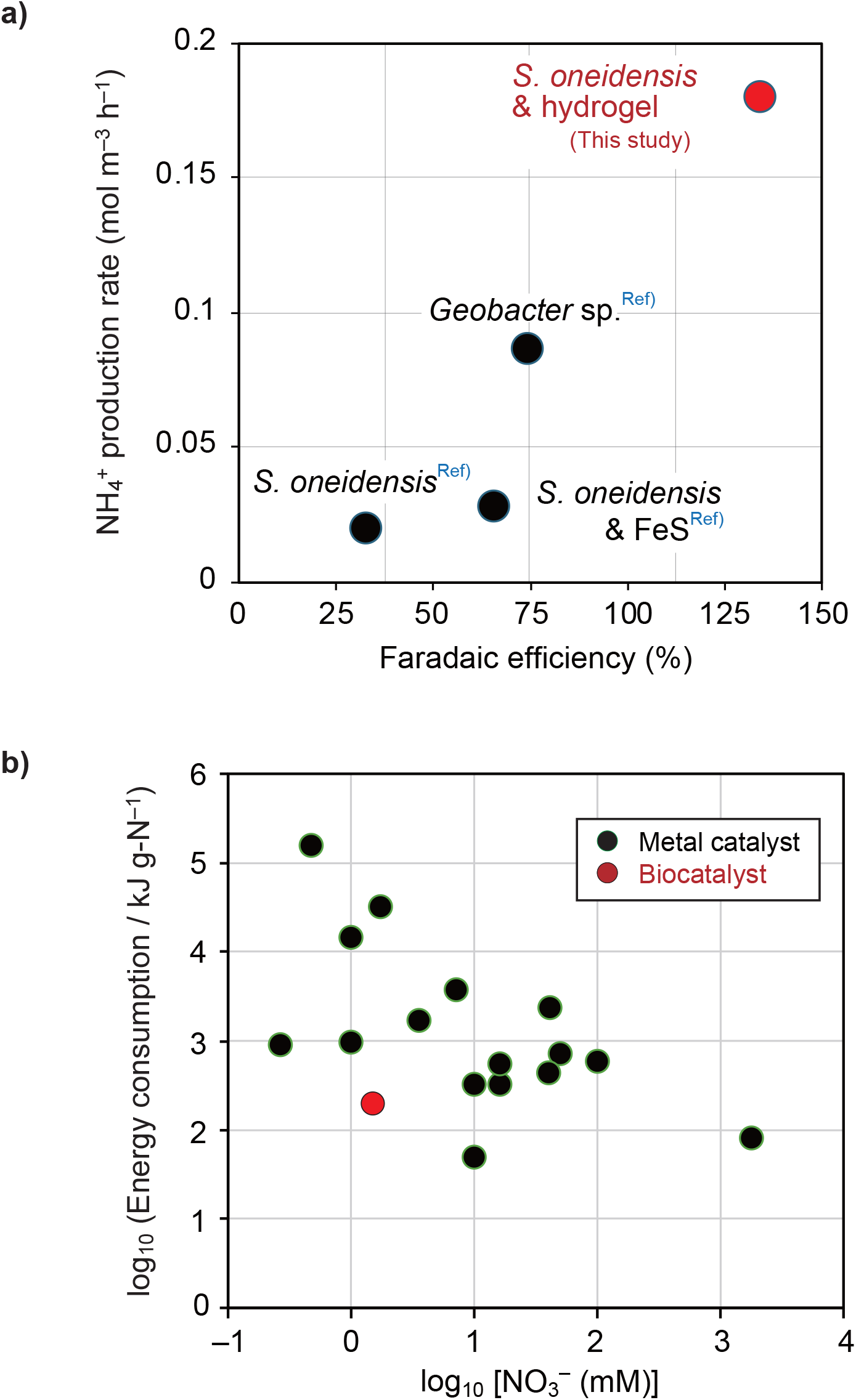
Performance comparison with previously reported systems. **(a)** NH_4_⁺ production rate vs. Faradaic efficiency (FE) for bioelectrochemical NO_3_⁻ reduction systems using biocatalysts. Data from previously reported systems (*Geobacter* sp., *S. oneidensis*, and *S. oneidensis* & FeS) are shown alongside the results of this study (*S. oneidensis* & hydrogel). **(b)** Relationship between initial NO_3_⁻ concentration (mM) and electricity consumption (kJ g-N⁻¹) for metal catalysts and biocatalysts reported in the literature, plotted on a log–log scale. Adapted from Kim *et al.* Nat. Commun. 14, 823 (2023), licensed under CC BY 4.0.

## 4. Discussion

### 4.1. Performance of bioelectrochemical NO_3_^-^ reduction to NH_4_^+^

The *S. oneidensis* hydrogel biocathode achieved the highest volumetric NH₄⁺ production rate among previously reported bioelectrochemical NO₃⁻ reduction systems using biocatalysts (Fig. 3a). Whereas conventional *S. oneidensis* biofilm-based systems required 3–5 days to reduce mM-level dilute NO₃⁻ [15,16], the hydrogel electrode in this study reduced the operational period to 0.5 days (Fig. 1b). Notably, the FE of the hydrogel electrode frequently exceeded the theoretical maximum of 100%, reaching values of 113–151% (Fig. 1e). Carry-over of organic carbon and nitrogen from the pre-culture LB medium into the cathode chamber was minimized by repeated washing of the cells and electrodes, and there are no reports indicating that the hydrogel components (CB/RF/SA/Cel) are assimilated as organic carbon sources by *Shewanella* [19]. Therefore, the FE values exceeding 100% are considered to reflect contributions from NH₄⁺ generation pathways other than the electrochemical reduction of NO₃⁻. Indeed, the ¹⁵N fraction of NH₄⁺ at the end of Cycle 1 (12 h) was 81 ± 1.9%, indicating that approximately 19% of the NH₄⁺ was generated as ¹⁴NH₄⁺. The most likely source of this ¹⁴NH₄⁺ is the mineralization of intracellular organic nitrogen compounds in *S. oneidensis* — including proteins, amino acids, nucleic acids, and cell wall constituents — released during microbial cell turnover (ammonification) [30]. The addition of this ¹⁴NH₄⁺ to the numerator of the FE calculation (total NH₄⁺ produced) is considered to have caused FE to exceed 100%.

Of particular interest, NO₂⁻ accumulation during NO₃⁻ reduction — frequently reported in previous *Shewanella* biofilm electrode studies [15,16], where up to approximately 1 mM NO₂⁻ accumulated during the reduction of 1.4 mM NO₃⁻ — was not observed in any cycle in the hydrogel system (Fig. 1b). Since NO₂⁻ is toxic to aquatic organisms and humans [31], its accumulation is undesirable. The absence of NO₂⁻ accumulation in the hydrogel system may be attributable to the possibility that electron supply to the ammonium-forming nitrite reductase (NrfA) was not rate-limiting in the hydrogel system, enabling NO₂⁻ to be consumed at a rate comparable to its production from NO₃⁻ reduction; alternatively, RF-mediated EET may have efficiently supplied electrons to both the nitrate reductase (Nap) and NrfA [32].

### 4.2 Mechanism of enhanced electron transfer in the hydrogel electrode

Since the amounts of *S. oneidensis* biomass on the hydrogel and biofilm electrodes was comparable, the higher NO₃⁻ reduction rate exhibited by the hydrogel electrode is interpreted as reflecting an improvement in electron transfer efficiency per unit *S. oneidensis* cell rather than an increase in cell abundance. Indeed, LSV showed that hydrogel modification increased the current density 6.1-fold (Fig. 2a), and this enhancement in current is considered to have directly contributed to the improvement in NH₄⁺ production rate. The correspondence between the increase in current and the increase in NO₃⁻ reduction rate suggests that the rate-limiting step in this system is the electron supply process from the cathode to the cells (EET), and that the hydrogel matrix plays a role in promoting this EET process. To gain deeper insight into the EET process, LSV was performed, revealing a distinct reduction peak at −0.648 V for the hydrogel electrode. This potential falls within the range of the redox potentials of flavins (including riboflavin) secreted by *Shewanella* and *c*-type cytochromes [33], suggesting that the RF incorporated into the hydrogel functions as an electron shuttle in mediated EET (MET) [34] between the membrane-bound cytochromes of *S. oneidensis* and the cathode surface. This interpretation is consistent with the findings of Yang *et al.* [19] in an anodic system, who demonstrated that adsorption of RF onto CB promotes MET and substantially reduces charge transfer resistance in a hydrogel bioanode. Also, the hydrogel electrode exhibited a markedly larger CV area compared to the biofilm electrode (Fig. 2b). The hydrogel electrode contains CB (Vulcan XC72R, BET surface area ≥200 m² g⁻¹), and the increased electrical double-layer capacitance attributable to its high specific surface area enables storage of a greater amount of charge [35]. This increase in effective electrode surface area is considered to increase the number of contact points between *Shewanella* and the electrode, thereby expanding opportunities for EET and contributing to the improvement in current density observed in LSV. On the other hand, our EIS measurements revealed no significant difference in charge transfer resistance (*R_ct_*) or solution resistance (*R_s_*) between the *S. oneidensis* hydrogel and biofilm electrode (Fig. 2c). This indicates that the higher current density of the hydrogel electrode does not result from a reduction in interfacial resistance but rather from RF–meditated MET that enables cells retained within the gel interior to contribute to electron transfer.

### 4.3 Stability of the hydrogel-based system across operational cycles

The hydrogel electrode maintained stable performance across Cycles 1 through 3, with no statistically significant decline observed in NH₄⁺ production rate, current, FE, or EC (Fig. 1c–f). In contrast, the biofilm electrode in which *S. oneidensis* was naturally attached to the electrode surface (Fig. S2) already exhibited markedly lower reduction rates than the hydrogel system in Cycle 1, and both NH₄⁺ production rate and current declined further in Cycle 2 following medium exchange. It is widely recognized that naturally formed biofilm electrodes are inherently susceptible to progressive performance deterioration caused by physical detachment and sloughing of the biofilm [36–38]. This limitation is particularly pronounced for *S. oneidensis*, which is known to form only thin biofilms as compared with *Geobacter sulfurreducens* (another electro active bacterium) [39,40], rendering the system vulnerable to biomass loss. The immobilization of *S. oneidensis* cells within an SA/Cel matrix overcomes this physical fragility and maintains stable performance over multiple cycles. This operational stability is an important characteristic for the application of this system to continuous or repeated water treatment processes.

Furthermore, the hydrogel electrode in this study was fabricated by applying the CB/RF/SA/Cel bioink to the graphite felt electrode via brush coating, a simple and low-cost method, without the use of a 3D bioprinter as employed in the preceding study by Yang et al. [19]. The results demonstrate that the electron transfer-enhancing effect of the CB/RF hydrogel — as reflected by the improvement in current density in LSV — can be reproduced by a simple coating operation without the need for high-precision manufacturing equipment such as 3D printers. This approach lowers the barrier to implementation and scale-up of the hydrogel electrode for practical water treatment processes. At the same time, there remains potential to further improve performance through the precise structural control offered by 3D printing, such as optimization of the microporous architecture within the gel and the spatial arrangement of cells and CB/RF [41], and a comparative investigation of the effect of fabrication method on electrode performance is warranted in future studies.

### 4.4 Comparison with metal catalysts and implications for dilute NO_3_^−^ treatment

Metal catalyst-based electrochemical NO₃⁻ reduction systems showed EC values on the order of 10^3^ kJ g-N⁻¹ at mM-level substrate concentrations (Fig. 3b). In contrast, the biocatalyst system in the present study achieved an EC of 1.68–2.39 × 10² kJ g-N⁻¹ at a comparable substrate concentration (initial NO₃⁻ concentration of approximately 1.5 mM), demonstrating substantially lower energy consumption than metal catalyst-based systems for the reduction of dilute NO₃⁻. This difference is attributable to two characteristics of biocatalysts compared to metal catalysts. First, biocatalysts can drive reactions at lower overpotentials [42]. Second, biocatalysts possess high apparent substrate affinity at low substrate concentrations; *e.g*., the apparent affinity constant (*K_m_*) of the nitrate reductase Nap; 3.4 µM for *Campylobacter jejuni* [43] and 120 µM for *Cupriavidus necator* H16 [44], and the *K_m_* for the NH_4_^+^ forming nitrite reductase Nrf; 54 µM for *S. oneidensis* MR-1 [45]. With respect to metal catalysts, it is well established that mass transfer limitations — specifically substrate diffusion limitations — become dominant under low-concentration conditions, necessitating greater overpotential (*i.e*., higher energy input) to maintain the reaction rate [46]. Under such conditions, the restricted migration of negatively charged NO₃⁻ ions toward the electrode surface and competition from the hydrogen evolution reaction further reduce energy efficiency. In contrast, biocatalysts possess enzymatic substrate recognition mechanisms characterized by high substrate affinity (low apparent *K_m_*), enabling efficient substrate capture and reduction even under dilute conditions, in accordance with Michaelis–Menten kinetics. These characteristics suggest that the present system may hold advantages over metal catalyst-based approaches for the treatment and resource recovery of dilute NO₃⁻ present in environmental waters at concentrations on the order of hundreds of µM to several mM.

### 4.5 Limitations and future directions

This study still has several limitations that need to be examined in other studies. First, this study employed a pure culture system of *S. oneidensis* MR-1, and the ability of *Shewanella* to persist and dominate in competition with the diverse indigenous microbial communities present in natural environmental waters has not been verified. In practical operation with natural waters, there is a possibility that *Shewanella* may be outcompeted by environmental microorganisms and displaced from within the hydrogel. It is therefore necessary to elucidate the long-term viability and competitive dominance of *S. oneidensis* within the hydrogel, as well as the maintenance of system performance in the presence of environmental microbial communities.

Second, the mechanistic interpretation of MET enhancement within the hydrogel proposed in this study is based on indirect evidence from electrochemical measurements including LSV, EIS, and CV, and does not account for the spatial arrangement of cells, RF, and CB within the hydrogel matrix. Future morphological observations of the spatial distribution of these components within the hydrogel — for example by SEM or CLSM — would enable more direct validation of the proposed mechanism.

Third, further improvement of the bioelectrochemical NO₃⁻ reduction rate is required for on-site treatment. Based on the surplus nitrogen of upland crops (approximately 36–203 kg-N ha⁻¹ y⁻¹) [47] and 21–46% of the nitrate leaching fraction of surplus nitrogen on arable land [48] (as cited in Fraters *et al*. [49]), the NO₃⁻ leaching rate from agricultural land is estimated at roughly 21– 256 g-N ha⁻¹ d⁻¹. Assuming on-site treatment with a cathode chamber volume of 1 m³ per hectare of agricultural land, the required NO₃⁻ reduction rate corresponds to 21–256 g-N m⁻³ d⁻¹ (0.063–0.76 mol m⁻³ h⁻¹). The previous and present bioelectrochemical systems achieve 0.02 – 0.19 mol m⁻³ h⁻¹ (Table 1), which is sufficient only for the least demanding case (*e.g*., sweet potato, 21 g-N m⁻³ d⁻¹), and for most upland crops the required rate exceeds the current performance, necessitating an improvement of up to approximately fourfold. For arable land on well-drained sandy soils, where the leaching fraction is higher (*i.e*, 80–99%) [49], the required rate rises further (up to ∼1.6 mol m⁻³ h⁻¹). Therefore, substantial further acceleration of the system is necessary to treat the NO₃⁻ loading from most agricultural land.

**Table 1.** Performance of bioelectrochemical NO_3_^-^ reduction to NH_4_^+^.

| Bacteria | Electrode | Electrode<br>size (cm<sup>3</sup>) | Reactor<br>volume (L) | Initial NO<sub>3</sub><sup>-</sup><br>(mM) | NH<sub>4</sub><sup>+</sup> production<br>rate (mol m<sup>-3</sup> h<sup>-1</sup>) | FE<br>(%) | Stability<br>(d) | Batch<br>Cycle | Ref. |
| --- | --- | --- | --- | --- | --- | --- | --- | --- | --- |
| *Geobacter*–<br>enriched<br>culture | Graphite rod | 4.7 | 0.1 | 9 | 8.6×10<sup>-2</sup> | 74 | < 5 | 1 | [17] |
| *S. oneidensis*<br>MR-1 | Carbon felt | 9.6 | 0.3 | 1.4 | 2.0×10<sup>-2</sup> | 32.7 | > 30 | 7 | [15] |
| *S. oneidensis*<br>MR-1@FeS | Carbon felt | 9.6 | 0.3 | 1.4 | 2.8×10<sup>-2</sup> | 65.4 | > 9 | 3 | [16] |
| *S. oneidensis*<br>MR-1 | CB/RF/SA<br>Hydrogel/<br>Graphite felt | 30 | 0.3 | 1.5 | 1.6 – 1.9 ×10<sup>-1</sup> | 113 – 151 | > 1.5 | 3 | This<br>study |

Finally, this study was limited to laboratory-scale validation (cathode chamber volume: 300 mL). Toward practical application to real environmental waters, future studies will need to address the effects of mass transfer limitations associated with scale-up to larger reactors, the long-term mechanical and biological stability of the hydrogel, and the cost of electrode materials.

## Conclusions

In the present study, the CB/RF/SA/Cel hydrogel electrode containing *S. oneidensis* MR-1 cells was fabricated, and its performance of bioelectrochemical NO₃⁻ reduction to NH₄⁺ was investigated under dilute NO₃⁻ conditions (*i.e*., 1.5 mM). The hydrogel electrode was prepared by the simple brush coating method, and maintained high (0.16 to 0.19 mol m^−3^ h^−1^) and stable (more than 1.5 d) NO₃⁻ reduction to NH₄⁺ performance across multiple cycles without NO_2_^−^ accumulation. Energy consumption for NH_4_^+^ production ranged from 168 to 239 kJ g-N^−1^. These findings demonstrate that the application of a hydrogel electrode represents a high-performance and stable approach for NH₄⁺ recovery from dilute NO₃⁻ via microbial electrosynthesis. Future work should address the long-term microbial community dynamics of this system in real environmental waters, further improvement of the NO₃⁻ reduction rate for on-site water treatment

## CRediT authorship contribution statement

Conceptualization: MO, SO

Investigation: RS, MO

Methodology: RS, MO

Formal Analysis: RS, YC, MO

Data curation: RS, YC, MO

Funding Acquisition: MO, SO

Supervision: MO, SO

Validation: MO, YC, SO

Visualization: MO, YC

Writing-original draft: MO

Writing-review & editing: MO, RS, YC, SO

## Data Availability Statement

The data of this study are available from the corresponding author upon reasonable request.

## Acknowledgements

This work was supported by JSPS KAKENHI (grant numbers 23H02114 and 25K01581 to M.O., 23H00192 to S.O.), JST FOREST Program (grant number JPMJFR216Z to M.O.), and JST NEXUS Program (grant number JPMJNX25F3 to M.O.). The authors sincerely thank Dr. Yasuhiro Kumaki (High-Resolution NMR Laboratory, Graduate School of Science, Hokkaido University) for assistance with NMR measurements, and Dr. Ryosuke Matsuo (Division of Environmental Engineering, Faculty of Engineering, Hokkaido University) for assistance with cyclic voltammetry measurements.

