## Supplementary file for "Hydrogel-enhanced bioelectrochemical nitrate reduction for ammonium recovery from dilute nitrate via S*hewanella oneidensis* MR-1"

**Supporting information**

This manuscript includes 2 supplementary figures and 7 supplementary tables.

**Fig. S1. Current profile during anodic biofilm formation on the carbon felt electrode.** *S. oneidensis* MR-1 was inoculated into the anode chamber with 18 mM lactate as the electron donor and the carbon felt electrode as the electron acceptor, and the electrode potential of the anode chamber was poised at +0.2 V vs. Ag/AgCl. Current increased following inoculation, reflecting microbial colonization and biofilm development on the electrode surface, and subsequently declined as lactate was depleted. The biofilm electrode formed after 60 h of operation was used as the biocathode in the subsequent $NO_3^-$ reduction experiment shown in **Fig. S2.**

**Fig. S2. Bioelectrochemical $NO_3^-$ reduction to $NH_4^+$ using a *Shewanella* biofilm electrode (without hydrogel). (a)** Time courses of $NO_3^-$, $NO_2^-$, and $NH_4^+$ concentrations (upper panels) and current (lower panels) during Cycle 1 (0–96 h) and Cycle 2 (0–216 h). **(b)** $NH_4^+$ production rate, **(c)** current (average value), **(d)** Faradaic efficiency (FE), and **(e)** electricity consumption (EC) for each cycle.

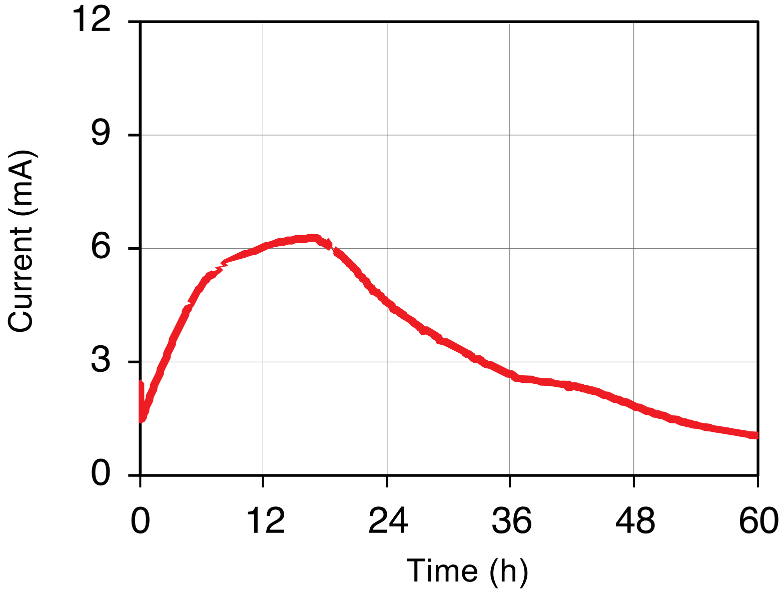


Fig. S1 (Oshiki *et al.*)

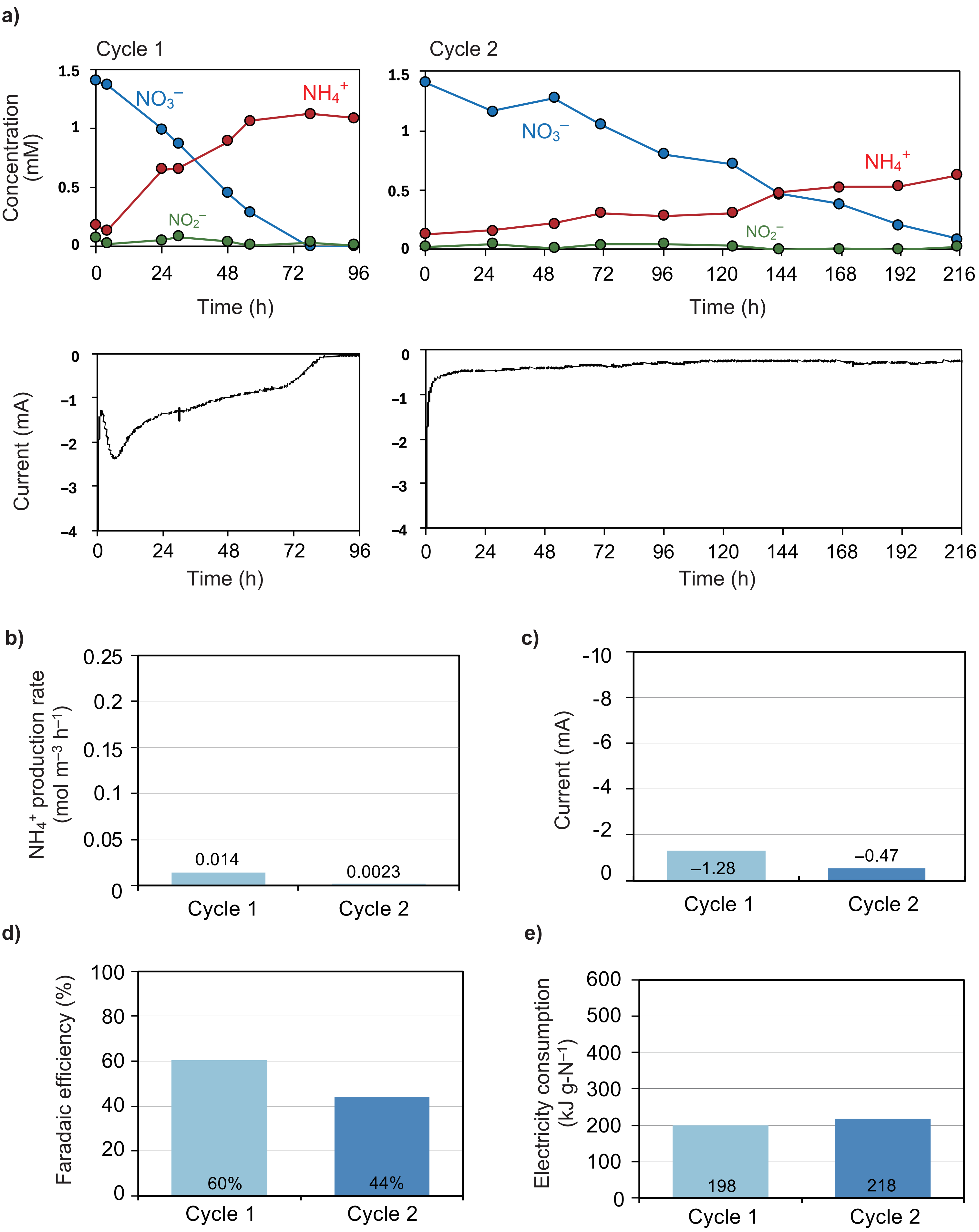


Fig. S2 (Oshiki *et al.*)

**Table S1 Anolyte composition for cultivating *S. oneidensis* MR-1 biofilm.**

| Components | Content |
|---|---|
| 10 × PBS (pH 7) | 100 mL/L |
| $C_3H_5NaO_3$ | 2.02 g/L |
| KCl | 0.13 g/L |
| $NaH_2PO_4$ | 2.56 g/L |
| $Na_2HPO_4$ | 4.09 g/L |
| $NH_4Cl$ | 0.31 g/L |
| mineral mix【**Table S2**】 | 12.5 mL/L |
| vitamin mix【**Table S3**】 | 5 mL/L |
| deionized water | 1,000 mL |

Preparation of media: Add components, except vitamin mix solution to deionized water, adjust pH 7.0 with 6 N NaOH or 6N HCl, and bring volume to 1,000 mL. Autoclave for 15 min at 15 psi pressure–120°C. Aseptically add 5 mL vitamin mix solution.

**Table S2 Composition of mineral mix.**

| Components | Content |
|---|---|
| Nitrilotriacetic acid | 1.5 g/L |
| $MgSO_4 \cdot 7H_2O$ | 3.0 g/L |
| $MnSO_4 \cdot 2H_2O$ | 0.5 g/L |
| NaCl | 1 g/L |
| $FeSO_4 \cdot 7H_2O$ | 0.1 g/L |
| $CoSO_4$ or $CoCl_2$ | 0.1 g/L |
| $CaCl_2 \cdot 2H_2O$ | 0.1 g/L |
| $ZnSO_4$ | 0.1 g/L |
| $CuSO_4 \cdot 5H_2O$ | 0.01 g/L |
| $AlK(SO_4)_2$ | 0.01 g/L |
| $Na_2MoO_4 \cdot 2H_2O$ | 0.01 g/L |
| $H_3BO_3$ | 0.01 g/L |

**Table S3 Composition of vitamin mix.**

| Components | Content |
|---|---|
| biotin | 2 mg/L |
| folic acid | 2 mg/L |
| pyridoxine hydrochloride | 10 mg/L |
| thiamine hydrochloride | 5 mg/L |
| nicotinic acid | 5 mg/L |
| DL-calcium pantothenate | 5 mg/L |
| vitamin $B_{12}$ | 0.1 mg/L |
| *p*-aminobenzoic acid | 5 mg/L |
| lipoic acid | 5 mg/L |

**Table S4 Catholyte composition for cultivating *S. oneidensis* MR-1 biofilm.**

| Components | Content |
|---|---|
| potassium ferricyanide | 50 mM |
| NaCl | 8.0 g/L |
| KCl | 0.2 g/L |
| $Na_2HPO_4$ | 1.42 g/L |
| $KH_2PO_4$ | 0.27 g/L |

**Table S5 Catholyte composition for bioelectrochemical $NO_3^-$ reduction**

| Components | Content |
|---|---|
| $NaNO_3$ | 1.4 mM |
| NaCl | 0.5 g/L |
| $NaH_2PO_4 \cdot 2H_2O$ | 5.04 g/L |
| $Na_2HPO_4 \cdot 12H_2O$ | 6.98 g/L |
| $NaHCO_3$ | 0.88 g/L |
| KCl | 0.7 g/L |
| Ca · Mg · Fe · EDTA mix 【Table S6】 | 250 mL |
| mineral mix【Table S2】 | 1 mL/L |
| vitamin mix【Table S3】 | 5 mL/L |
| deionized water | 750 mL |

Preparation of media: Add components, except Ca · Mg · Fe · EDTA mix, vitamin mix solution to deionized water, adjust pH 6.7 with 6 N NaOH or 6N HCl, and bring volume to 750 mL. Autoclave for 15 min at 15 psi pressure–120°C. Aseptically add 250 mL Ca · Mg · Fe · EDTA mix, and 5 mL vitamin mix solution.

**Table S6 Composition of Ca · Mg · Fe · EDTA mix.**

| Components | Content |
|---|---|
| $MgSO_4 \cdot 7H_2O$ | 0.41 g/L |
| $CaCl_2$ | 0.02 g/L |
| $FeSO_4 \cdot 7H_2O$ | 0.07 g/L |
| EDTA-2Na | 0.07 g/L |
| deionized water | 250 mL |

Preparation of media: Add the components to deionized water and bring volume to 250mL. Autoclave for 15 min at 15 psi pressure–120°C.

**Table S7 Anolyte composition for bioelectrochemical $NO_3^-$ reduction.**

| Components | Content |
|---|---|
| NaCl | 8.00 g/L |
| KCl | 0.2 g/L |
| $Na_2HPO_4$ | 1.42 g/L |
| $KH_2PO_4$ | 0.27 g/L |
| deionized water | 1,000 mL |
